# MTD: a unique pipeline for host and meta-transcriptome joint and integrative analyses of RNA-seq data

**DOI:** 10.1101/2021.11.16.468881

**Authors:** Fei Wu, Yao-Zhong Liu, Binhua Ling

## Abstract

RNA-seq data contains not only host transcriptomes but also non-host information that comprises transcripts from active microbiota in the host cells. Therefore, joint and integrative analyses of both host and meta-transcriptome can reveal gene expression of microbial community in a given sample as well as the correlative and interactive dynamics of host response to the microbiome. However, there are no convenient tools that can systemically analyze host-microbiota interactions by simultaneously quantifying host and meta-transcriptome in the same sample at the tissue and the single-cell level, which poses a challenge for interested researchers with a limited expertise in bioinformatics. Here, we developed a software pipeline that can comprehensively and synergistically analyze and correlate the host and meta-transcriptome in a single sample using bulk and single-cell RNA-seq data. This pipeline, named MTD, can extensively identify and quantify microbiome, including viruses, bacteria, protozoa, fungi, plasmids, and vectors in the host cells and correlate the microbiome with the host transcriptome. MTD is easy to install and run, involving only a few lines of simple commands. It empowers researchers with unique genomics insights into host immune responses to microorganisms.

## 1 Introduction

A variety of microorganisms have been recognized to contribute to the development of human diseases, including cancer, autoimmune diseases, and psychological disorders. For example, Helicobacter pylori can cause stomach cancer [1], human papillomavirus (HPV) infection can lead to uterine cervix cancer [2], and HIV-1 infection [3] can lead to HIV-Associated Neurocognitive Disorders (HANDs) [4]. In addition, the Epstein–Barr virus (EBV) was found to contribute to 1.5% of total human cancers of various types worldwide [5]. Therefore, systemic and comprehensive investigation of microorganisms in tissues for their contribution to disease development is critical, and in particular, the pathogenic mechanisms of opportunistic infections need to be fully dissected. Furthermore, it is necessary to address differential abundance of distinct cell populations as it may contribute to the heterogeneous tropism in infections. Therefore, it is important to analyze microbiome diversity, abundance, their interaction with host cells, and impacts on infected cells at both bulk tissue and single cell level.

Transcriptome from a host tissue may contain mRNAs from microorganisms that have not been fully investigated. Several tools have been developed to detect the microbiomes in the RNA-seq data, such as Kraken2 [6], VIRTUS [7], and IDseq [8]. However, there is no convenient tool that can analyze both host and microbiome transcriptome in the same set of data within a single workflow of analyses, which poses a challenge to researchers, especially those without bioinformatics expertise, who are interested in examining microbiome in host tissues and its relation to the endogenous expression of host genes at the both bulk tissue and single-cell level. To facilitate such effort of joint analyses of host transcriptome with its microbiome, a user-friendly pipeline, MTD (Meta-Transcriptome Detector), were developed for comprehensive and integrative investigation of microbiome from bulk and single-cell RNA-seq data.

## 2 Methods

### 2.1 Description of MTD

The MTD has two sub-pipelines to detect and quantify microbiomes by analyzing bulk and single-cell RNA-seq data, respectively (Figure 1). MTD is written in R (version 4.0.3) and Bash (version 4.2) languages and executed in GNU/Linux system. Users can easily install and run MTD using only one command and without requiring root privileges. The outputs (graphs, tables, count matrixes, etc.) are automatically generated and stored in the designated directory/folder defined by the user. The user manual for detailed instruction of installation and usage is available on the webpage https://github.com/FEI38750/MTD. Here we describe the two sub-pipelines separately.

**Figure 1.**
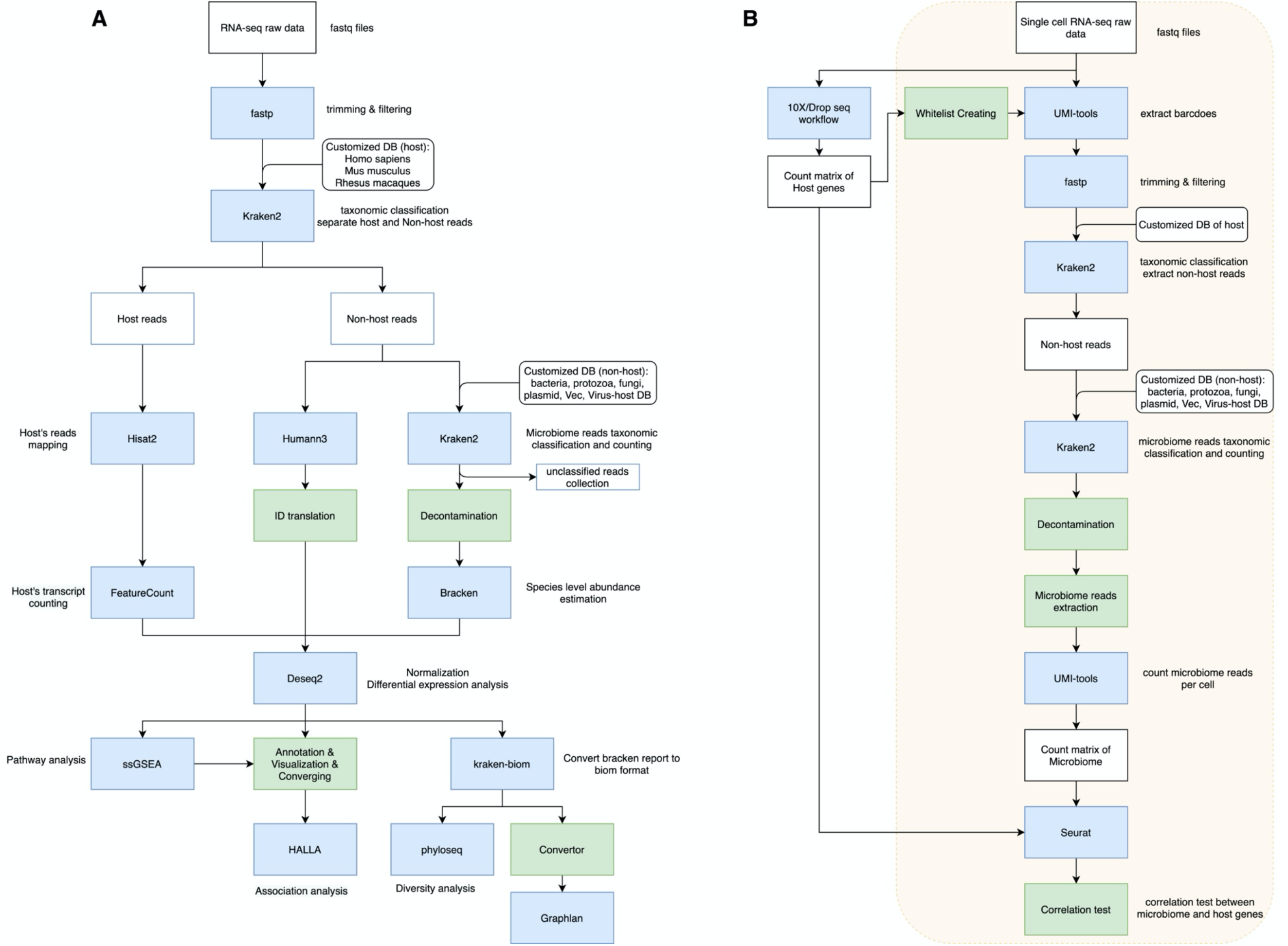
An overview of MTD. (A): A workflow for bulk mRNA-seq analysis. (B): A workflow for single-cell mRNA-seq analysis. White boxes represent the reads in FASTQ format and the count matrix. Blue boxes show the bioinformatics software used. Green boxes are the additional tools for data processing. The white boxes with curved edges show the reference genome and databases. In the single-cell mRNA-seq workflow (B), the left side exemplifies the host reads process protocols, and the right side in yellow shadow shows the MTD automatic pipeline to calculate the count matrix for the microbiome reads and the correlation test between microbiome and host genes.

#### 2.1.1 Transcriptome and meta-transcriptome analysis of bulk RNA-seq data

First, RNA-seq raw reads in the FASTQ file are trimmed and filtered by fastp (version 0.20.1) [9] with polyA/T trimming, and reads shorter than 40 bp (with the option --trim_poly_x -- length_required 40) are discarded. Then, processed reads are classified based on the host genome by Kraken2 (version 2.1.1) [6] with default parameters. Finally, the host and non-host reads are organized separately in FASTQ format.

##### Host transcriptome analysis

By default, the host species supported by MTD are Homo sapiens (Reference genome assembly: GRCh38), Mus musculus (Reference genome assembly: GRCm39), and Macaca mulatta (Reference genome assembly: Mmul_10). However, if needed, users can add other host species by one command line (bash Customized_host.sh -t [threads] -d [host_genome_Ensembl_address] -c [host_taxid] -g [host_gtf_Ensembl_address]). The host reads are aligned to the reference host genome by Hisat2 (version 2.2.1) [10] with default parameters written in a SAM file. Quantification of reads for host gene expression is done by featureCounts (version 2.0.1) [11]. Next, the count data is analyzed by DESeq2 package (version 1.32.0) [12] in Bioconductor to obtain the differentially expressed genes (DEGs). The gene annotation is done through the biomaRt R package (version 2.46.3) [13, 14] in Bioconductor. The data visualization and a count matrix are automatically generated through R programs. The data visualization includes the heatmap, Venn Diagram, PCA, barplot, and volcano plot. The count matrix contains the Ensembl gene ID, gene symbol, chromosome name, gene position, functional descriptions, DEG results for each pairwise group comparison, raw, normalized and transformed reads counts. This count matrix is saved in CSV format, ready for downstream analyses such as pathway enrichment and customized data visualization.

##### Microbiome transcriptome analysis

MTD supports a broad spectrum of microbiome species and vectors including viruses, bacteria, protozoa, fungi, plasmids, and vectors. At the time of writing, the viruses contain 16,275 species from Virus-Host DB [15] and also simian immunodeficiency virus (SIVmac239) (GenBank accession number M33262). The rest of the microbiome are from the NCBI RefSeq database [16], which includes 63,237 species of bacteria, 13,970 fungi, 1,337 archaea, 573 protozoa, and 5,855 plasmids. In addition, vector contamination can be screened using the NCBI UniVec Database. Users can update the microbiome databases in MTD by one command line (bash Update.sh -t [threads]). The non-host reads are further classified by Kraken2 based on microbiome references with the default parameters, followed by a decontamination step that removes the microorganism under the genera reported as reagent and laboratory contaminants [17]. Users can customize the blacklist of contaminants based on their experimental environments. Then, the abundance of the microbiome in the species level is calculated by Bracken (version 2.6.0) [18]. Next, the count data is imported into DESeq2 for analysis of the differentially expressed species. In DESeq2, we adjust the abundance of microbiome species based on the transcriptome library size of the sample. The rationale is that counts from a microbial species should take into account the overall representation of the host transcriptome. This normalization step is conducted through a formula in DESeq2, *design ~ group + transcriptome_size*, where transcriptome_size is defined by the formula: log2 (of a transcriptome size) - mean (of all log2-transformed transcriptome sizes in a sample), and group is the group code (e.g., A or B) of a sample. As a result, the importance of a microbial species in a sample was adjusted for the overall representation of host transcriptome. Meanwhile, the non-host reads (including both unclassified and classified reads by Kraken2 using the reference databases) are imported into the Humann3 (version 3.0) [19] for profiling microbial metabolic pathways and molecular functions. The ChocoPhlAn [20] and full UniRef90 [21] are used as reference databases for nucleotide and protein, respectively. Then the profiling results are annotated to commonly used functional terms to facilitate the downstream analyses. Next, the heatmaps of DEGs, Venn Diagrams, PCA, bar plots, and volcano plots are generated for the microbial species, molecules, and metabolic pathways based on the Deseq2 results. Additionally, kraken-biom (version 1.0.1) [22] is used to format the data for diversity analyses and phylogenetic tree plotting. The phyloseq R package (version 1.34.0) [23] in Bioconductor and vegan R package (version 2.5-7) [24] are used to analyze the diversities, including alpha diversity (Shannon, Simpson) and beta diversity (Bray-Curtis). Then, box plots are shown to visualize the t-test comparison results between groups, including alpha diversities of classified reads and the abundances of unclassified reads. PCoA graphing and analysis of similarities (ANOSIM) are based on Bray-Curtis distance. The relative abundance of a microbial species in the total microbiome is shown as bar plots at the Phylum level. The heatmap of total microbiome abundance is plotted using data normalized by Deseq2. Next, the phylogenetic trees are plotted through modified Graphlan [25], which is a tool for generating informative and integrative circular graph representing phylogenetic and taxonomic trees. However, the original program requires a specific data format as input and is not compatible with the output from Kraken2 and Bracken software. Therefore, we wrote a converter program and integrated it into the pipeline to bridge the taxonomic classification software Kraken2/Bracken and graph-making software GraPhlAn. It allows us to transform the output of Kraken2 and Bracken to match the data structure requirement of GraPhlAn. Furthermore, the default settings of colors have been optimized by modifying the source code of GraPhlAn. In addition to .biom format, data is also saved in .mpa and .krona formats to facilitate further downstream visualizations.

Most importantly, in the final steps of the pipeline, MTD examines the association between the microbiome and the host’s characteristics, such as gene expression and pathways. Pathway enrichment for each sample is performed by the single sample Gene Set Enrichment Analysis (ssGSEA 2.0) program [26–28] with a MSigDB C2 database that contains 6,290 curated gene sets. ssGSEA 2.0 is modified for parallel computing in High-Performance Computing (HPC) environment, which is described in the supplementary document. Next, the effects from covariates among groups are adjusted through the removeBatchEffect function in the limma R package (version 3.48.1) [29] in Bioconductor. The association analyses are then conducted through Halla (version 0.8.18) [30], which is set to compute hierarchical clustering of Spearman pairwise correlation. Figure 2 illustrates the MTD automatic pipeline for the bulk RNA-seq raw data.

**Figure 2.**
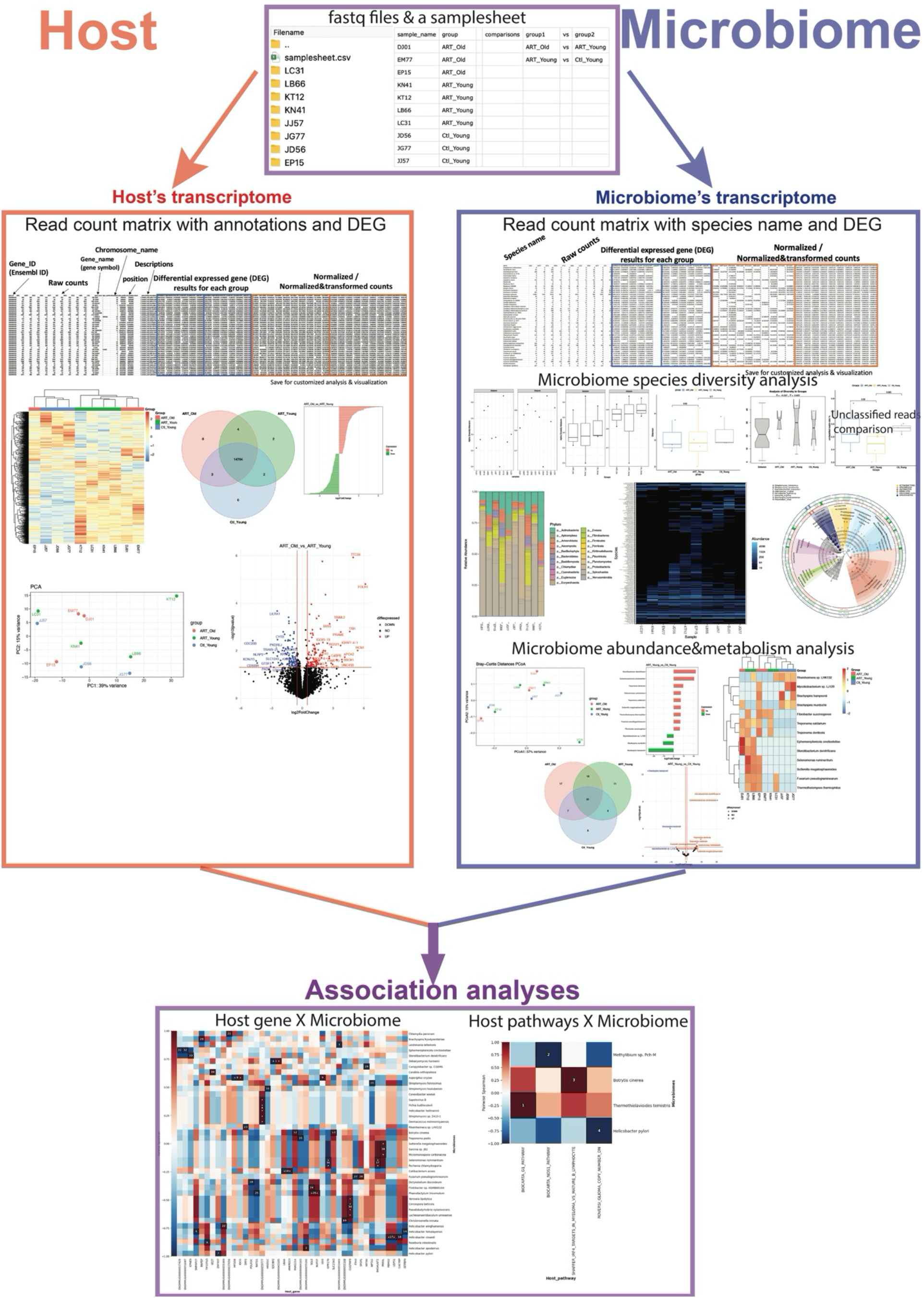
The automatic pipeline of MTD for analysis of the bulk RNA-seq raw data. Analysis results are automatically saved into the folder assigned by the user. Examples of analysis outputs for the transcriptome of host and microbiome are demonstrated on the left side and right side, separately. The shared procedures are shown in the purple boxes, which include the input files (upper box) and the association analyses (bottom box). In addition to the graphs, all the detailed information was included in the data sheets and stored in the corresponding output folder. To use the MTD, the user needs to place the FASTQ files in a folder with a sample sheet in CSV format that describes the sample names, groups, and comparisons, and then perform the analysis with one command line. For example, with the command line “bash MTD.sh -i~/inputpath/samplesheet.csv -o ~/outputpath -h 9544 -t 20”, the user provides the place of the sample sheet with the raw data after the flag “-i”, and the location to save the results (after flag “-o”), the host taxonomic ID is after “-h”, and the threads of CPU after “-t”.

#### 2.1.2 Analysis of single-cell RNA-seq data

The MTD supports automatic generation of the count matrix of the microbiome by using raw data in the FASTQ format and count matrix of host genes from two commonly used single-cell RNA-seq platforms, 10x Genomics and Drop-seq. The user may start with a count matrix of host genes (H5, Matrix, or .dge.txt format) or follow the 10x Cell Ranger or Drop-seq pipeline if starting with the raw data. The cell barcodes are identified from the count matrix of host genes, then the UMI and cell barcodes are extracted and added to read names by using the UMI-tools (version 1.1.1) [31]. The reads are then trimmed and filtered by fastp, and filtered for the host reads by Kraken2. The non-host reads are further classified by Kraken2 with the comprehensive microbiome databases, followed by a decontamination step. The steps from trimming to decontamination use the same settings as mentioned in the previous section. Next, the taxonomic labels of the reads are extracted and aligned with corresponding cell barcodes through a step written by the AWK program language. Finally, UMI-tools are used to generate a count matrix (tab-separated) through a converter written in R. Figure 3 exemplifies a count matrix automatically generated by MTD for the microbiome in each cell. Subsequently, MTD combines the count matrices of the host genes and microbiome to perform the correlation analysis automatically. An example of the correlation analysis result is showed in Figure 7 (A-B).

**Figure 3.**
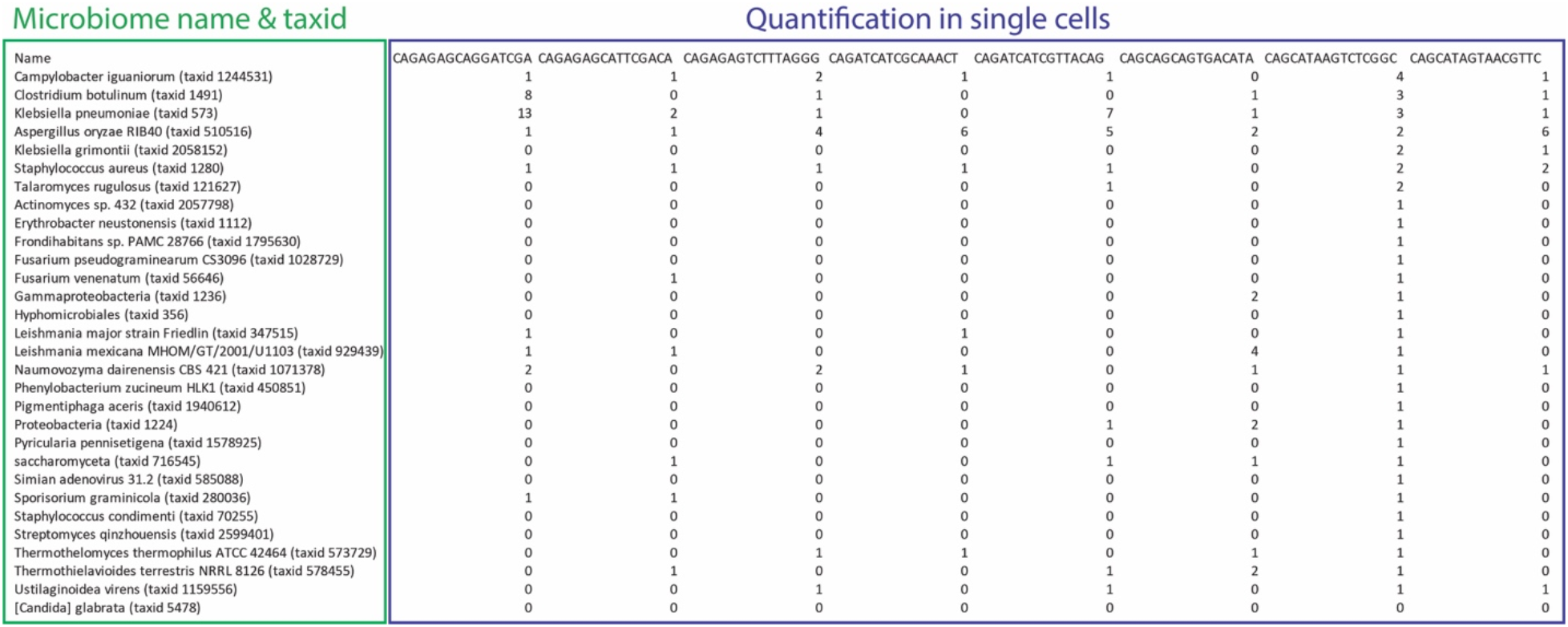
An example of the count matrix that is automatically generated by MTD for the single-cell microbiome analysis. For illustration, the figure shows part of the large count matrix. The name and taxonomy ID of the microbiome is shown in the first column and highlighted in the green box. The read counts are highlighted in the blue box. The first row shows the cell barcodes.

At the single-cell level, the Spearman correlations between microbial organisms and host genes are tested by using the top 3,000 most highly variable features, including the normalized data of both host and microbiome. Because the step of correlation analysis step is highly time-consuming for a large data matrix, parallelizing computing was applied to speed up the computation by using the doParallel R package (Version 1.0.16). The other analysis methods through Seurat R package (Version 4.0.1) and homemade programs are described in the supplementary document. The diagram of the MTD pipeline for single-cell RNA-seq is demonstrated in Figure1 B.

### 2.2 Animal information and sample collection

Here we demonstrated the application of MTD for analyzing the bulk RNA-seq data by using samples from the descending colon from rhesus monkeys as an example. The detailed sample information is in the supplementary document.

Raw data of the singe-cell RNA-seq was downloaded from NCBI GEO with accession code GSE161340 (SRR13041553-13041560), and GSE160384 (SRR12933210-SRR12933217). The detailed sample preparation methods for the sequencing were described in the articles [32, 33]. The detailed methods for single-cell RNA-seq analysis are in the description in supplementary document.

## 3 Results

### 3.1 Bulk RNA-seq analysis: descending colon of rhesus macaques

Through MTD, the transcripts of both the microbiome and the host were analyzed simultaneously using the same bulk RNA-seq raw data. Figure 4 is a heatmap showing the abundance of all the microbiome species in the samples from the descending colon. Supplementary Figure 1 presents the taxonomic and phylogenetic trees of microorganisms detected in the descending colon samples from rhesus macaques. More detailed results of the microbiome and host gene analyses of the descending colon are described in the supplementary document sections 2.1 and 2.2, respectively. The microbiome analysis results of BMC are in supplementary document sections 2.3. Here we demonstrate the results of association analyses of the microbiome to the host genes or pathways in descending colon.

**Figure 4.**
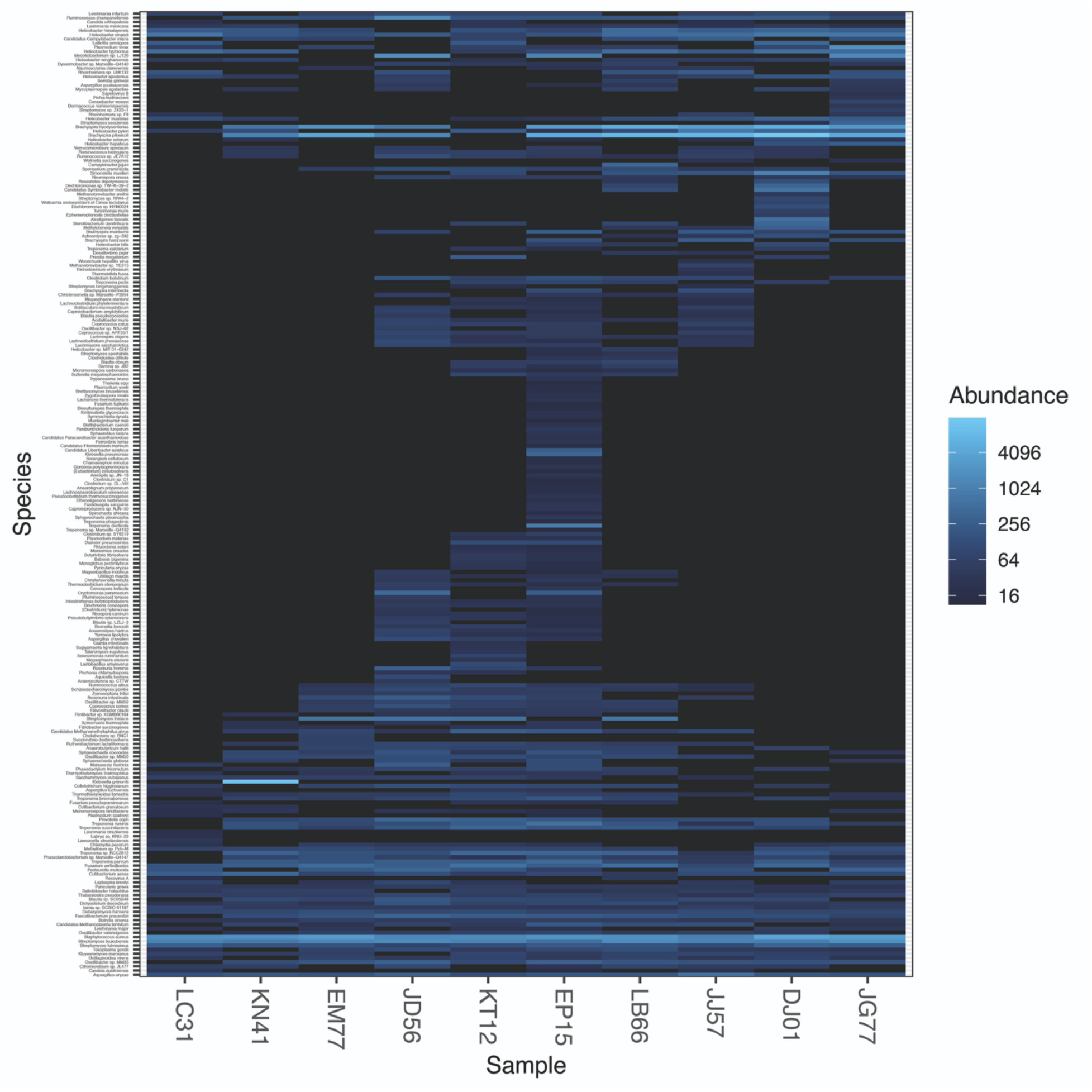
Heatmap of the microbiome species identified in descending colon samples of rhesus macaques. The higher abundance of microbiome reads is shown in a light blue color, and the lower abundance is shown in a darker color. Data was normalized using the Deseq2 and plotted by the phyloseq R packages, which were wrapped in MTD.

The correlations between the microbiome and host gene are illustrated in Figure 5A. For example, the *Debaryomyces hansenii* shows a significant positive correlation with host gene *UBE2I* and *IL27RA*. The expression of the host gene *C1QTNF8* is positively correlated with a group of microbes, such as *Yarrowia lipolytica*, *Aspergillus chevalieri*, *Roseburia hominis*, and *Anaerostipes hadrus*.

**Figure 5.**
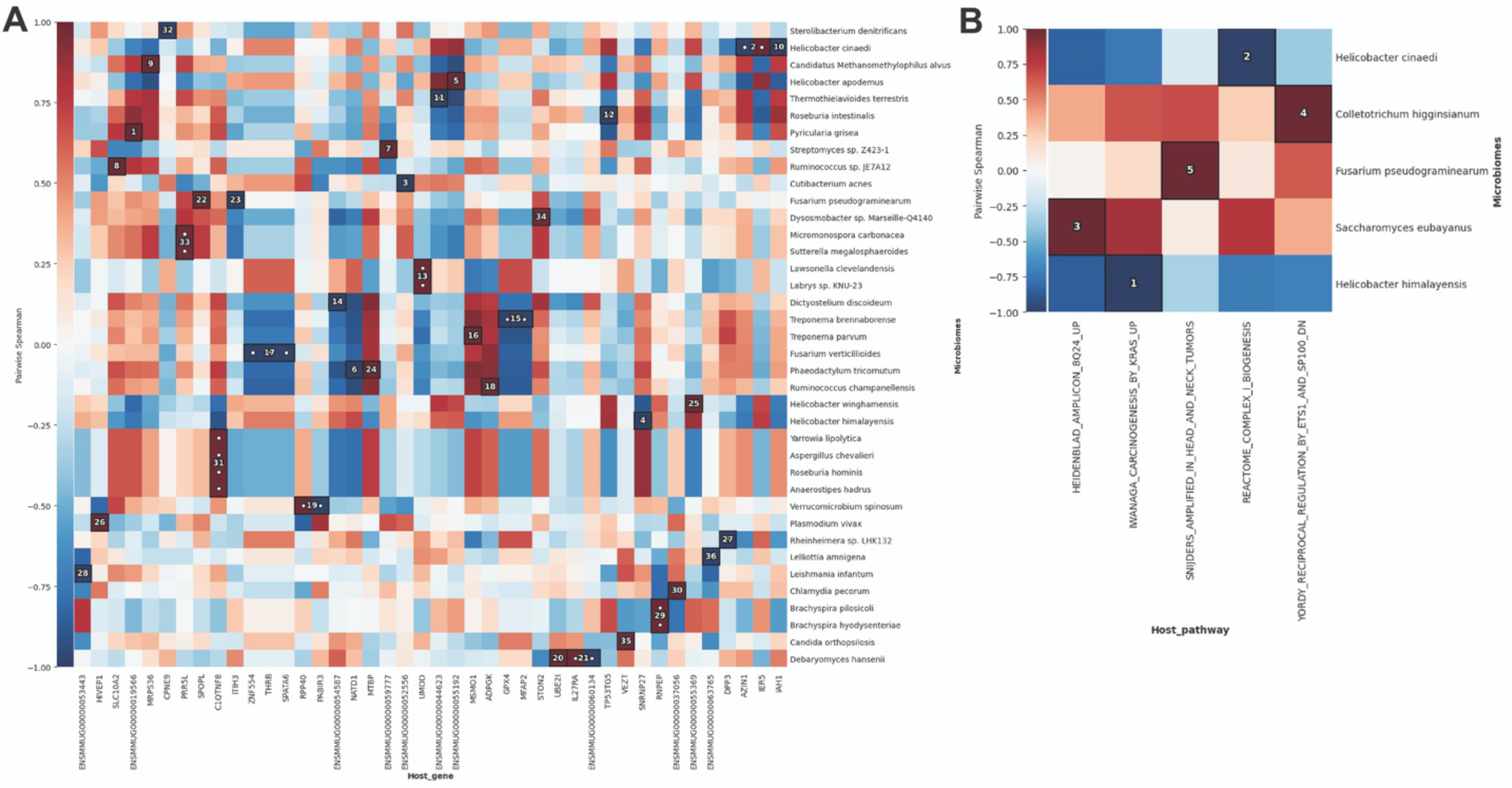
Analyses of association between the microbiome and host genes or pathways in descending colon of the rhesus macaque. The figure shows the correlation between the RNA expression level of microbiome species and host gene (A), or pathways (B). The x-axis is labeled with the names of the host genes or pathways, and the y-axis lists the names of microbiome species. Positive correlation coefficients are shown in red, and negative correlation coefficients are shown in blue. The significant results are marked with white dots and ranked by numbers. The results in the same cluster can be found in a box with the same number. The association was examined by pairwise Spearman correlation test. The tabulates of all comparison results and dot plots were saved in the corresponding output folder.

The correlations between the microbiome and host pathways are shown in Figure 5B. For example, the pathway that controls the amplification of the 8q24 chromosome region (HEIDENBLAD_AMPLICON_8Q24_UP) was upregulated with the mRNA expression of *Saccharomyces eubayanus*. The complex I biogenesis signaling pathway (REACTOME_COMPLEX_I_BIOGENESIS) was negatively correlated with the expression of *Helicobacter cinaedi*.

### 3.2 Single-cell RNA-seq analysis

#### 3.2.1 Microglia cells of SIV-infected rhesus macaques

We next applied MTD to single-cell RNA-seq data from microglia cells isolated from SIV-infected rhesus macaques [32]. Because the analysis results from the authors identified SIV transcripts in the single cells, it is an ideal dataset for validation of the capacity of MTD to process single-cell RNA-seq data.

First, count matrices of the microbiome were generated by MTD. Then, they were integrated with the host transcriptome for downstream analysis through the Seurat R package. The results showed that the major cell type in the sample was microglia, with a small portion of endothelial cells (Figure 6A). A cluster of microglia cells was highly infected with SIV (Figure 6B). For this cluster, the top 20 makers ranked by fold change are displayed in Figure 6C, such as PDE4A and SENP3. The cell subpopulation with these markers implicated a higher SIV tropism.

**Figure 6.**
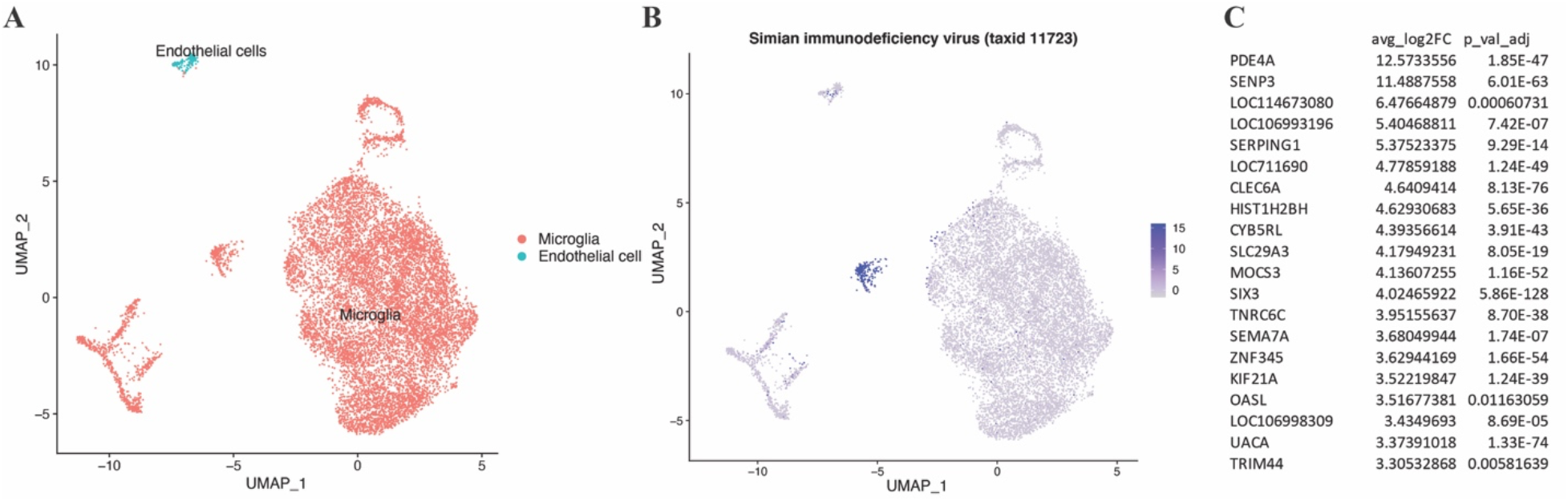
Detection of SIV in microglia cells from rhesus macaques. (A) Cell types in different colors on the UMAP plot. Cell type identity was assigned based on the homemade program described in supplementary document. (B) SIV reads detected by MTD. The blue dots on the UMAP plot indicate the SIV-infected cells with the normalized reads quantity. (C) Markers of the cell cluster that harbor SIV. Analyses of the count matrix followed by visualization were performed through Seurat. FindMarkers function with MAST methodology was used for computing the log2fold changes for each gene/feature between clusters and their corresponding adjusted p-values.

Overall, the results validated the capability of MTD for detecting specific microorganism species from single-cell RNA-seq data.

#### 3.3.2 Brain cells of mice

We also applied MTD on the single-cell RNA-seq data of brain cells isolated from mice [33] and demonstrated the correlation analyses between the microbiome and host genes or pathways at the single-cell level.

We found that *plasmodium vivax* (*P*. *vivax*) and host gene *GRIA2* had the highest correlation coefficient (Figure 7A). As shown in Figure 7B, *GRIA2* was highly expressed in the *P. vivax*-infected cells. We further identified all the host genes that highly correlated with *P. vivax* (Figure 7C), then performed pathway enrichment analysis. The results underscored the positive association of *P. vivax* with the function of the infected cell’s plasma membrane region, such as cell junction and transmembrane transporter activity (Figure 7D). This result supports the cytoadherence phenomenon of *P. vivax* reported in previous research [34-36]. Although *P. vivax* primary infects red blood cells, its cytoadherence on other cell types has been reported, such as in endothelial cells [34, 35]. Moreover, recent findings suggest that it has the ability to adhere to all Chinese hamster ovary (CHO) cells [35]. Our results brought insights into the interaction between *P. vivax* and host cells. It showed that *P. vivax* interacts with host cells that are incrementally expressing genes of the cellular membrane. Future research can study the causal effect of these molecules during infection, such as whether they contribute to pathogen adherence or if the infection leads to their increased expression.

**Figure 7.**
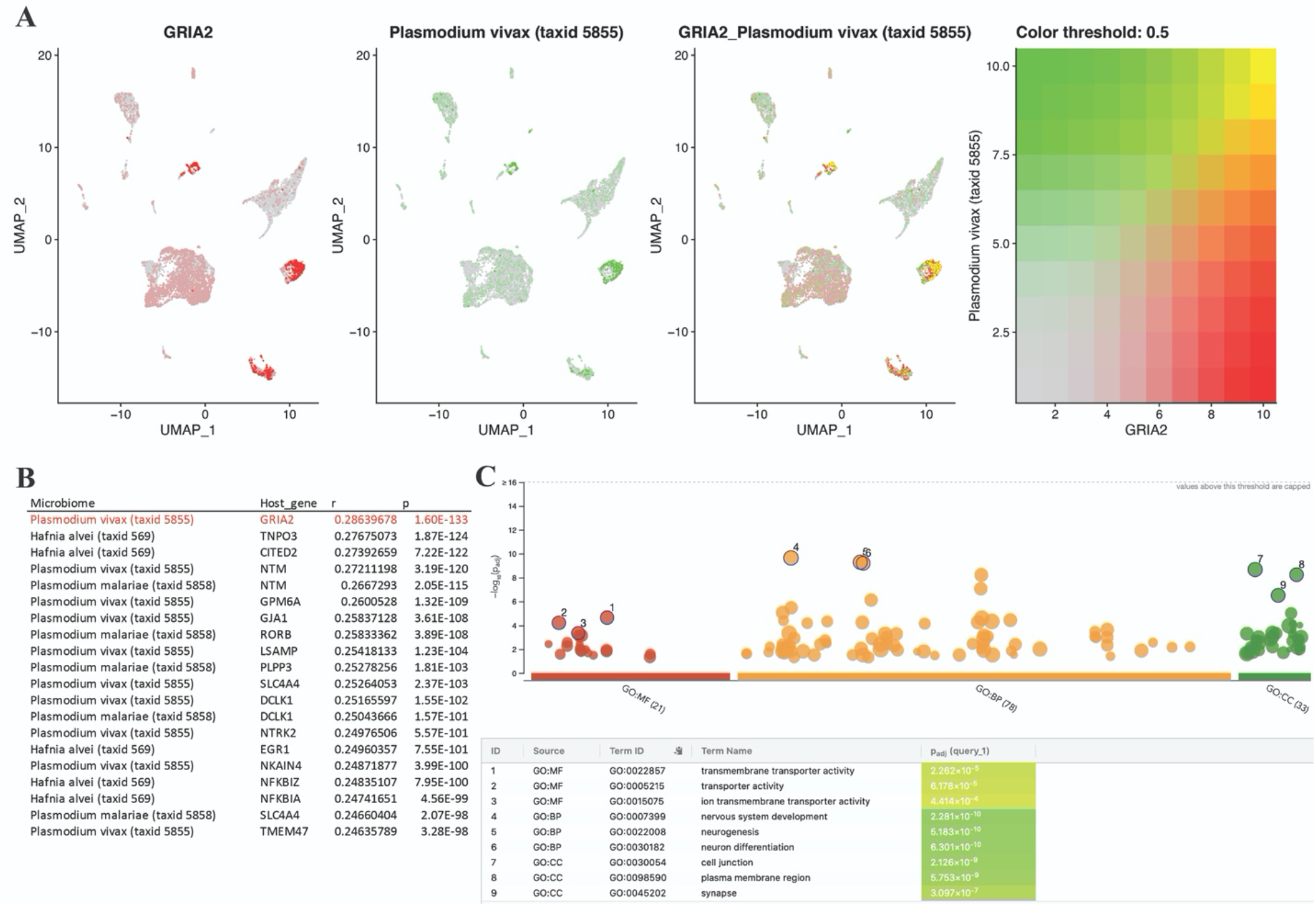
Co-expression of microbiome and host genes in host cells. (A) Visualization of the highest correlation on the UMAP plot. The cells expressed *GRIA2* are represented by red dots, and the cells infected with *plasmodium vivax* are shown as green dots. The cells containing both are represented by the overlapping of the two colors (yellow). (B) A list of the top 20 correlations between host genes and microorganisms, and the highest pair is highlighted in red. (C) The results of pathway enrichment of the genes that were highly associated with *plasmodium vivax*, which were defined by r >0.2 and p < 0.05. The top 3 results of each GO categories are highlighted. UMAP plots were drawn by the Seurat R package. The pathway enrichment was performed through g:GOSt in g:Profiler (version: e104_eg51_p15_3922dba, organism: mmusculus).

## 4 Conclusion

MTD is a powerful and convenient host and meta-transcriptome joint and integrative analyses pipeline for both bulk and single-cell RNA-seq data. With the use of this software pipeline, the activated microbiome (including the virome) can be detected, the cell type harboring them can be identified, and correlation between the host gene expression and microbial prevalence can be directly analyzed. Thus, it would be a powerful tool to improve our understanding of host-pathogen interactions, in particular, how the microbiome contributes to the host health and disease, and what genes and pathways of the host are important to a particular infection by a microbial species, which may shed lights to prevention and treatment of common human diseases from a metagenomics perspective.

## 5 Discussion

MTD has several unique advantages compared to current tools in detecting meta-transcriptome. MTD can simultaneously and comprehensively profile microbiome and the host transcriptome in both bulk and single-cell RNA-seq data. The program takes into account the host transcriptome library size while modeling microbiome reads in differential abundance analysis. In addition, the program has a decontamination step to eliminate the potential noise from the contaminant microbes, including the common contaminant microbes in the laboratory environment [17]. Users can also modify the list of contaminant microbes to better suit their specific requirements. MTD warrants easy usability as well as data safety. The software installation, updating, and analyses can be achieved by single lines of command. It obviates the need of using other cloud-based applications.

Currently, it is common to perform the polyA tail enrichment during the library preparation for mRNA sequencing. Thus, meta-transcriptomics analyses as implemented with MTD could avoid contamination from viruses in most cases because virus RNA only acquires polyA tail when it is transcribed in the host cell. However, users need to be cautious about the possibilities of single-strand RNA viruses and other preparation methods that contain the polyA tail. Nevertheless, it is challenging to detect and remove other contaminated microbes. There are tools to identify the potential contaminant by simply calculating the correlation of nucleic acid abundance between microbes and host [37]. However, these methods may not be sensitive if the samples have similar abundance, heterogeneous contaminant patterns, or cross-contamination.

In the era of single-cell genomics, the presence of the microbiome in the cell populations and their associations with cell functions need to be better analyzed and further understood. With MTD, researchers are now able to acquire genomics insights into the pathogenesis of microorganisms identified. Furthermore, annotating the sample’s geographic information with each microorganism would offer us a map of pathogens, which could predict an epidemic. Thus, MTD could also become a critical element for monitoring the spread of the microbiome and its pathogenesis in the future.

### Key Points

- MTD enables simultaneous analyses of the microbiome and the host cell transcriptome in bulk and single-cell RNA-seq data.
- The correlation between the microbiome and the host transcriptome can be automatically analyzed.
- MTD has an extensive microbiome detection capacity, including viruses, bacteria, protozoa, fungi, plasmids, and vectors.
- Installation and use MTD is as easy as one command line without the requirement of administrator/root privilege.
- Decontamination function is enabled to eliminate the common contaminant microbes in the laboratory environment.

## Supplementary Data

Supplementary data are available online at *Briefings in Bioinformatics*.

## Data Availability

MTD software can be accessed through GitHub at https://github.com/FEI38750/MTD

## Funding

This work was supported by National Institutes of Health R01 MH116844 (BL) and R01 NS104016 (BL). The funders had no role in study design, data collection and analysis, preparation of the manuscript or decision for publication.

## Acknowledgements

The authors thank Sandra Smith and Brian Kopecki at the Department of Information Technology of Texas Biomedical Research Institute to maintain HPC and install the third-party programs asked. We are grateful to any contributors in our respectable open-source community and their virtue in sharing the questions and answers.

## Notes

https://github.com/FEI38750/MTD

### Competing Interest Statement

The authors have declared no competing interest.

### Summary of Updates

Adjusted the wording, rephrased some sentences, and rearranged the supplementary document examples to make them clearer.

