## Supplementary Document for "MTD: a unique pipeline for host and meta-transcriptome joint and integrative analyses of RNA-seq data"

### 1 Methods

#### 1.1 Animal information

For bulk RNA-seq of descending colons, a total of 9 Chinese rhesus macaques were used, including 6 SIV-infected and ART-treated and 3 SIV-uninfected controls (Supplementary table 1).

Animal euthanasia was performed in line with the recommendations of the Panel on Euthanasia of the American Veterinary Medical Association. Following Tulane IACUC standards of operation (SOP), Simian immunodeficiency virus (SIV)-infected and/or drug-treated macaques were euthanized first with telazol and buprenorphine, followed by sodium pentobarbital intravenous injection. Tissues from descending colon were collected fresh and soaked in RNAlater™ Stabilization Solution during necropsies. To avoid the chance of microbiome contamination from the animal gut to the brain, brain tissues were collected first, and the descending colons were collected last.

#### 1.2 Bulk RNA-seq

RNA was prepared using RNeasy Plus Mini Kit (QIAGEN Cat#74134). mRNA sequencing experiment was performed by Novogene. The PolyA selected and non-strand specific library was prepared by NEBNext® UltraTM RNA Library Prep Kit for Illumina®. About 20 million paired-end 150 bp reads per sample were generated from Illumina NovaSeq 6000 Sequencing System.

The RNA-seq raw reads stored in the FASTQ format were analyzed by MTD. Rhesus macaque’s genome (Mmul_10 assembly) and its annotation (Macaca_mulatta.Mmul_10.103.gtf) from the Ensembl genome database were used for reads alignment and counting. Significant DEG were defined by log2FoldChange > 0.5 and p-value < 0.05.

#### 1.3 Single cell RNA-seq analysis

Raw data of the singe-cell RNA-seq was downloaded from NCBI GEO with accession codes GSE161340 (SRR13041553-13041560) and GSE160384 (SRR12933210-SRR12933217) by using the SRA Toolkit. The detailed sample preparation method for the sequencing was described in the articles [1, 2].

Reads type in SRA was first checked by the command like vdb-dump SRR13041553 -R1 -C READ_TYPE then converted to the raw FASTQ files by fasterq-dump function (e.g., fasterq-dump SRR13041553 --include-technical -S) in SRA Toolkit. Downstream processes were conducted on the corresponding biological and technical reads in FASTQ files.

Count matrix of the single-cell microbiome was generated through MTD software. In R, the count matrixes of the microbiome and host from the same sample were preprocessed then merged by the rbind function. Downstream analysis was performed with the Seurat R package (Version 4.0.1) [3-6]. In Seurat, the datasets were first normalized with the SCTransform function, then followed the integration workflow to joint analysis of multiple single-cell datasets.

SCTransform method is more effective than a traditional normalization workflow and eliminates more technical effects from data. It allows higher PCs, which make confounding factor that due to variation in sequencing depth is substantially mitigated. Consequently, higher PCs represent more heterogeneity that is both subtle and biologically relevant - so including them during downstream analysis can potentially improve results. Moreover, this method returns more variable features by default, which may represent more subtle biological changes [7].

Data under "SCT" assay of the Seurat object (e.g. project_name@assays[["SCT"]]@data) was log-normalized using SCTransform function. The top 3,000 highly variable genes/features among the datasets were used to find "anchors" for integration. UMAP projections were calculated for the first 30 principal components of the log-normalized data. UMAP was chosen instead of the t-SNE because it could provide a better separation of identified cell types [8]. FindMarkers function with MAST methodology was used for computing the log-fold changes for each gene/feature between groups and their corresponding p- and adjusted p-values [9].

A homemade cell type classifier was used to identify the senescent cells and other canonical cell types in the brain. It was achieved by performing the single sample Gene Set Enrichment Analysis (ssGSEA) [10-12] for each cell based on the gene expression profile of distinct cell types. A list of senescence marker genes was referred from a recent single-cell analysis study on mice brains [13]. Marker gene lists of cell types were downloaded from PanglaoDB database [14]. Brain and vasculature were selected as organs contain cell types present in the brain.

A modified ssGSEA2.0 [11] program was used for the enrichment score calculation. High Performance Computing (HPC) cluster/server had been used to accelerate the calculation. Because ssGSEA2.0 demands almost all (maximum threads -1) CPU processing capacity for its multi-threads computation and without provides related settings, which makes its usage on the HPC cluster/server troublesome. This greedy occupation of the computational resource of the server would result in fail the job submission, disturb other users, or be banned by the administrator. So, in order to make the program more compatible with the server, we have modified it to allow users to easily adjust the demands of threads to an appropriate number based on their actual computational environment. Then, normalized enrichment scores (NES) were calculated for every cell within the data set based on the degree of absolute enrichment of a gene set.

The attributes of cells (e.g., cellular senescence group, senescent score, age group, viral infection status, cell types) were annotated and stored in the metadata of the Seurat object (e.g., project_name@meta.data). The virus abundance in the cell was indicated by the virus reads count ratio, which was calculated by dividing the number of all of the reads that belong to the virus by the total reads number in the cell. The Pearson correlations between cell types were examined by combine using the cor (for correlation coefficient), cor.test (for p-value), and corrplot (for plot) functions in Base R and corrplot R packages. The Pearson correlations between senescent cells and virus abundance were calculated and plotted by ggscatter function in ggpubr R package. The source code of analyses are in the supplementary material GSE160384_analysis, CellTypes_Ident, and GSE161340_analysis.

### 2 Results

#### 2.1 Microbiome analysis of descending colon

An overview of differentially expressed microbiome species are shown in Supplementary Figure 2A, and the relative abundances of microbiomes at the phylum level were plotted in Supplementary Figure 2B. The heatmap with the detailed name of each differentially expressed species is stored in the same output folder defined by the user. The clusters groups were plotted on PCoA based on Bray-Curtis distance (Supplementary Figure 2C). Three groups were not well separated in the PCoA, and the analysis of diversity did not find a significant difference among the groups (Supplementary Figure 2D). A hundred and seven species were shared by these groups, and ART_Old animals had more unique species than other groups (Supplementary Figure 2E). The reads that did not belong to the host were also examined, including all the microbiome information, classified and unclassified (Supplementary Figure 2F). ART young animals showed a relatively lower non-host reads ratio than other groups, especially compared to the nonSIV control young animals, which implicated the effect of ART in reducing microbial abundance. However, the alpha-diversity was not significantly different among groups (Supplementary Figure 2G). In general, the results indicated a lack of disparities in the microbiome diversity in the descending colons between the groups of ART_Old, ART_Young, and Ctl_Young. More detailed information about the significantly differentially expressed microbiome species between groups can be found in Supplementary Figure 3.

The results of metabolic molecules analyses are shown in Supplementary Figure 4-5. In addition, just a few microbial metabolic pathways were profiled in our samples. Among them, aerobic respiration I (cytochrome c) is the only significantly differently expressed pathway, which was higher in the Ctl_Young group than the other two groups. The table of pathway analysis results was stored in the corresponding output folder.

#### 2.2 Host gene expression analysis of descending colon

ART_Young and Ctl_Young showed more dissimilarities in host gene expression patterns, whereas ART_Old cannot be separated from the other two groups (Supplementary Figure 6 A-B). Most of the genes were shared among the three groups (Supplementary Figure 6 C). The overview of significant DEG between groups are shown in the thumbnail bar plots (Supplementary Figure 6 D, F) and the volcano plots (Supplementary Figure 6 E, G). The top 20 genes with the most significant absolute fold changes are labeled in the volcano plots.

In addition, the comprehensive count matrix, the DEG table of each comparison, and the heatmap with gene names were stored in the corresponding output folder defined by the user. They can easily be used for downstream analyses, such as pathway enrichment, functional annotation, and molecular interaction network.

### Discussion

RNA-seq data includes both host and non-host information. For the RNA-seq method, the microbiome must infect the cells and utilize the host’s cellular system to express its mRNA, which also means the activate microbiome in the host cells. These conditions differ from the microbiota in fecal samples, where the microbiome is not necessary to infect the host cells. Therefore, this transcriptome analysis of the active microorganism in the host cell provides further direct information about pathogen-host interactions.

MTD has several advantages compared to current tools in detecting meta-transcriptome.

1. MTD enables simultaneous detection of the microbiome (including the virome) in the cells and the host gene expression in both bulk and single-cell RNA-seq data. To our knowledge, current other tools have limited microbiome species coverage (partial bacteria or viruses) or are unable to identify the microbiome in single-cell RNA-seq data. This is partially due to the conventional alignment-based methods that have significantly high hardware requirements and slow processing speeds for detecting an extensive range of microbiomes. MTD uses an alignment-free detection approach, which is much faster and has relatively low hardware requirements.
2. Owing to the alignment-free detection approach, MTD is resistant to low sequence conservation due to mutation and recombination events [38] that often occur in microorganisms.
3. MTD has a better accessibility. Many current analysis tools depend on Docker, or require administrator/root privilege to install, which restricts their usage on most scientific computing resources, such as HPC with multi-user systems. MTD does not require administrator/root privilege for users to install and use on HPC. Moreover, some other tools can only be accessed through specific user accounts [8, 39], further limiting accessibility, especially for users not supported by NIH funding [39].
4. MTD warrants better data safety. MTD can be easily deployed on a local HPC of an institute or laboratory. Users do not need to apply particular accounts and upload their data to an outside cloud-based software.
5. MTD takes into account the host transcriptome size while modeling microbiome reads in differential abundance analysis, which better reflects the infectious severity of the exogenous species on the host.
6. MTD analyzes the association between the microbiome and the host gene expression or pathways. Thus, this information could offer researchers valuable insights into pathogen-host interaction.
7. MTD has an extensive virome detection ability. Comprehensive virome information from the Virus-Host database was included in the MTD. In addition to the NCBI RefSeq and GeneBank, it also contains references from UniProt, Viral Zone, and manually curated annotations.
8. MTD uses the two-step approach to gain better specificity [7]. First, host reads are extracted, then the microbiome is classified in the remaining non-host reads. Thus, through the alignment-free approach, host reads could be identified and removed more explicitly, which further minimizes contamination from host reads during the microbiome analysis.
9. MTD is a user-friendly software that can be executed by one command line. It minimizes the programming knowledge requirements for its usage. Moreover, the classified reads in the intermediate steps are saved in the FASTQ format, which is beneficial for users who need additional analyses.
10. MTD has flexible host species choice. It initially supports three commonly used host species: human, mouse, and rhesus monkey. In addition, users can add other host species by just one command line.
