## Supplementary Figure 1 for "MTD: a unique pipeline for host and meta-transcriptome joint and integrative analyses of RNA-seq data"

### Slide 1
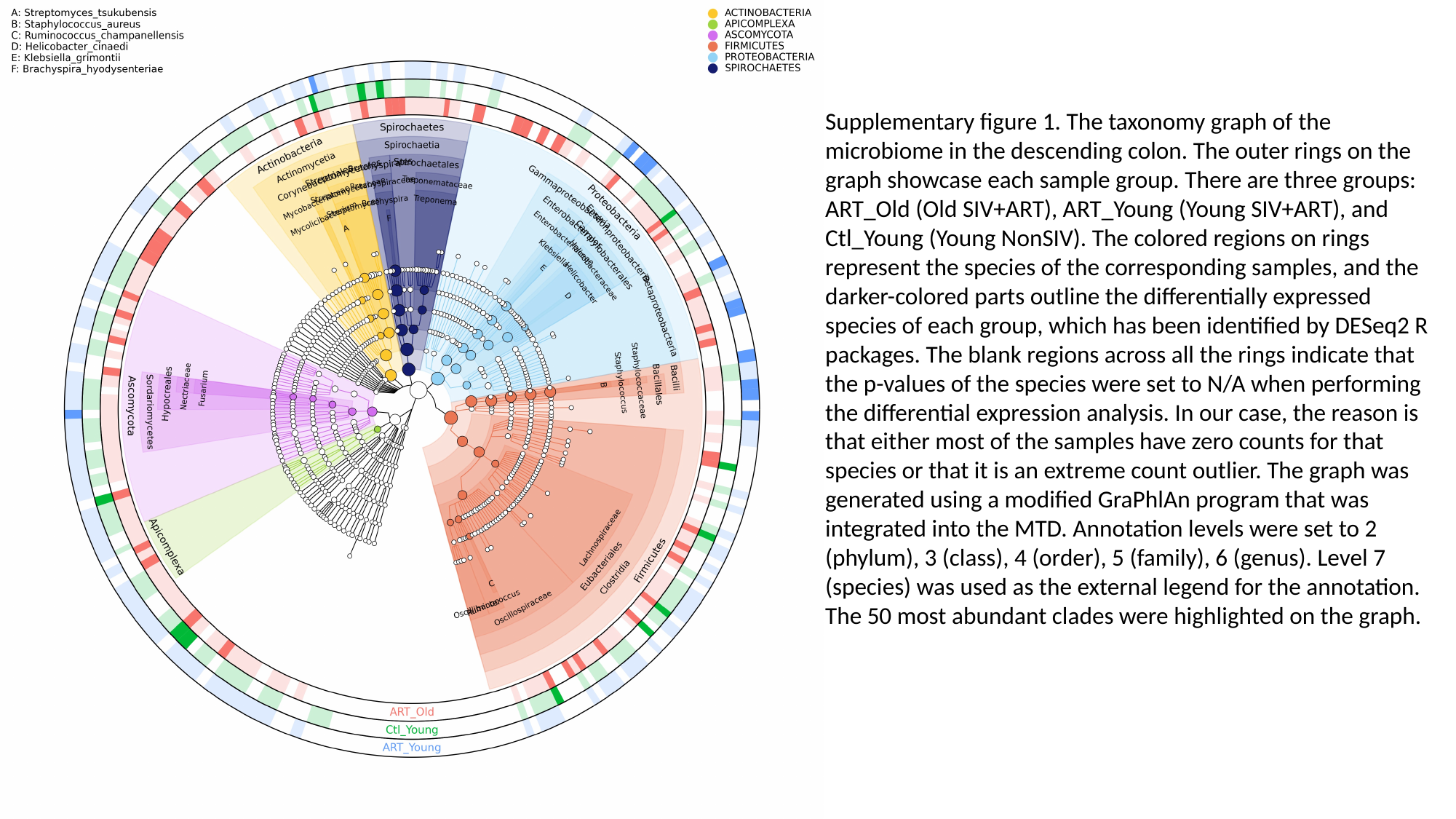

Supplementary figure 1. The taxonomy graph of the microbiome in the descending colon. The outer rings on the graph showcase each sample group. There are three groups: ART_Old (Old SIV+ART), ART_Young (Young SIV+ART), and Ctl_Young (Young NonSIV). The colored regions on rings represent the species of the corresponding samples, and the darker-colored parts outline the differentially expressed species of each group, which has been identified by DESeq2 R packages. The blank regions across all the rings indicate that the p-values of the species were set to N/A when performing the differential expression analysis. In our case, the reason is that either most of the samples have zero counts for that species or that it is an extreme count outlier. The graph was generated using a modified GraPhlAn program that was integrated into the MTD. Annotation levels were set to 2 (phylum), 3 (class), 4 (order), 5 (family), 6 (genus). Level 7 (species) was used as the external legend for the annotation. The 50 most abundant clades were highlighted on the graph.
