## Supplementary Figure 2-3 for "MTD: a unique pipeline for host and meta-transcriptome joint and integrative analyses of RNA-seq data"

### Slide 1
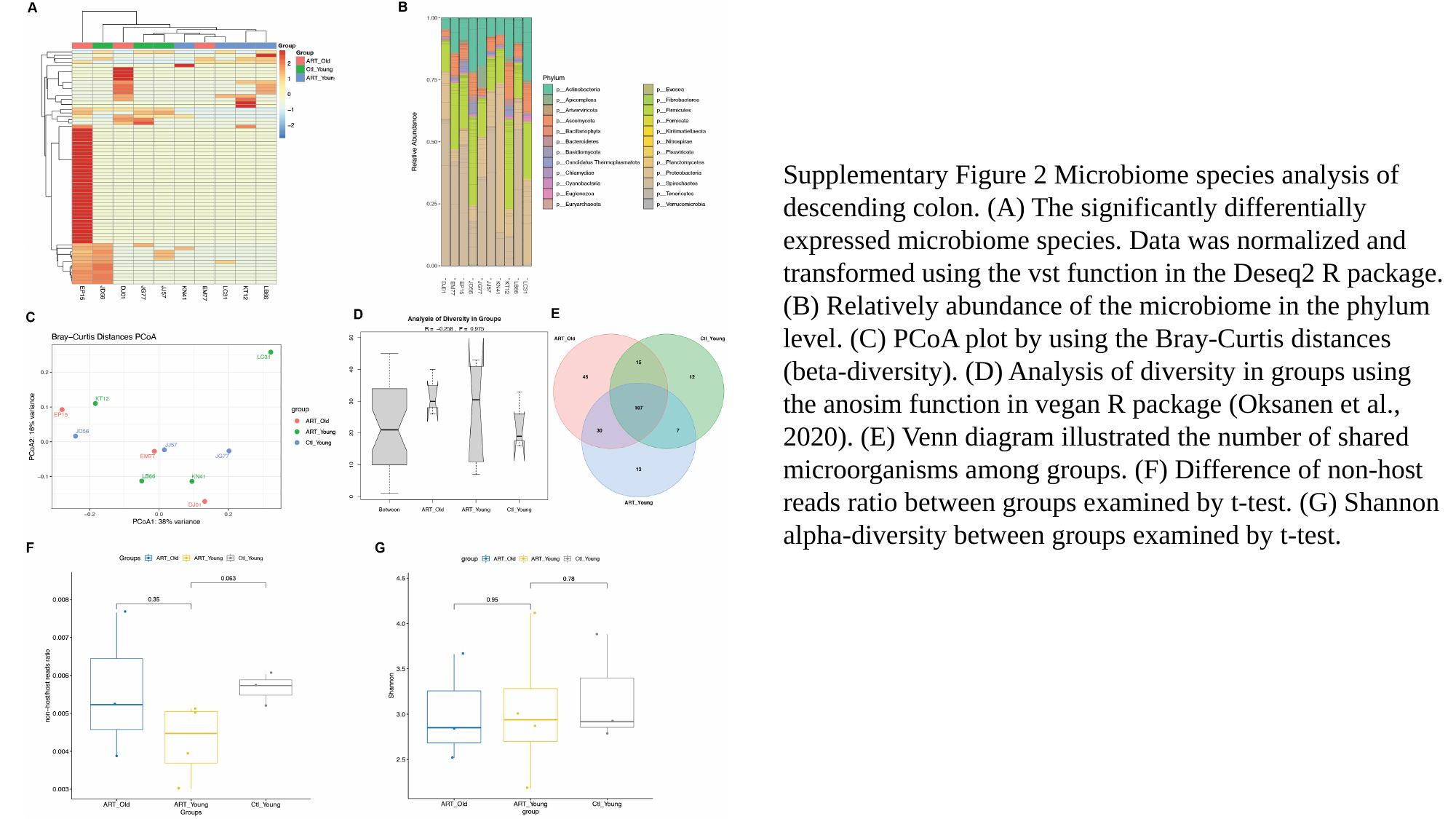

Supplementary Figure 2 Microbiome species analysis of descending colon. (A) The significantly differentially expressed microbiome species. Data was normalized and transformed using the vst function in the Deseq2 R package. (B) Relatively abundance of the microbiome in the phylum level. (C) PCoA plot by using the Bray-Curtis distances (beta-diversity). (D) Analysis of diversity in groups using the anosim function in vegan R package (Oksanen et al., 2020). (E) Venn diagram illustrated the number of shared microorganisms among groups. (F) Difference of non-host reads ratio between groups examined by t-test. (G) Shannon alpha-diversity between groups examined by t-test.

### Slide 2
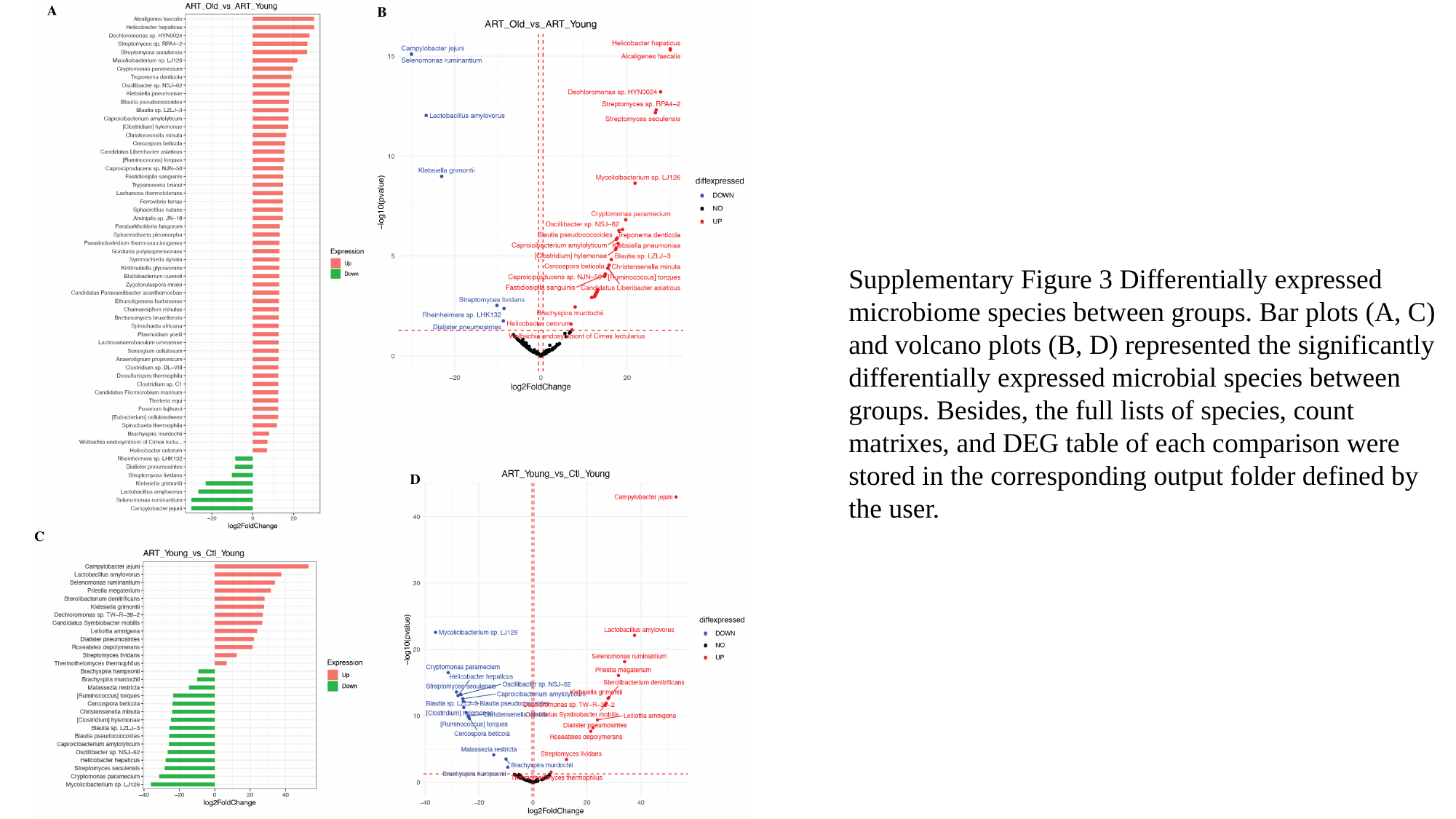

Supplementary Figure 3 Differentially expressed microbiome species between groups. Bar plots (A, C) and volcano plots (B, D) represented the significantly differentially expressed microbial species between groups. Besides, the full lists of species, count matrixes, and DEG table of each comparison were stored in the corresponding output folder defined by the user.
