## Supplementary Figure 4-5 for "MTD: a unique pipeline for host and meta-transcriptome joint and integrative analyses of RNA-seq data"

### Slide 1
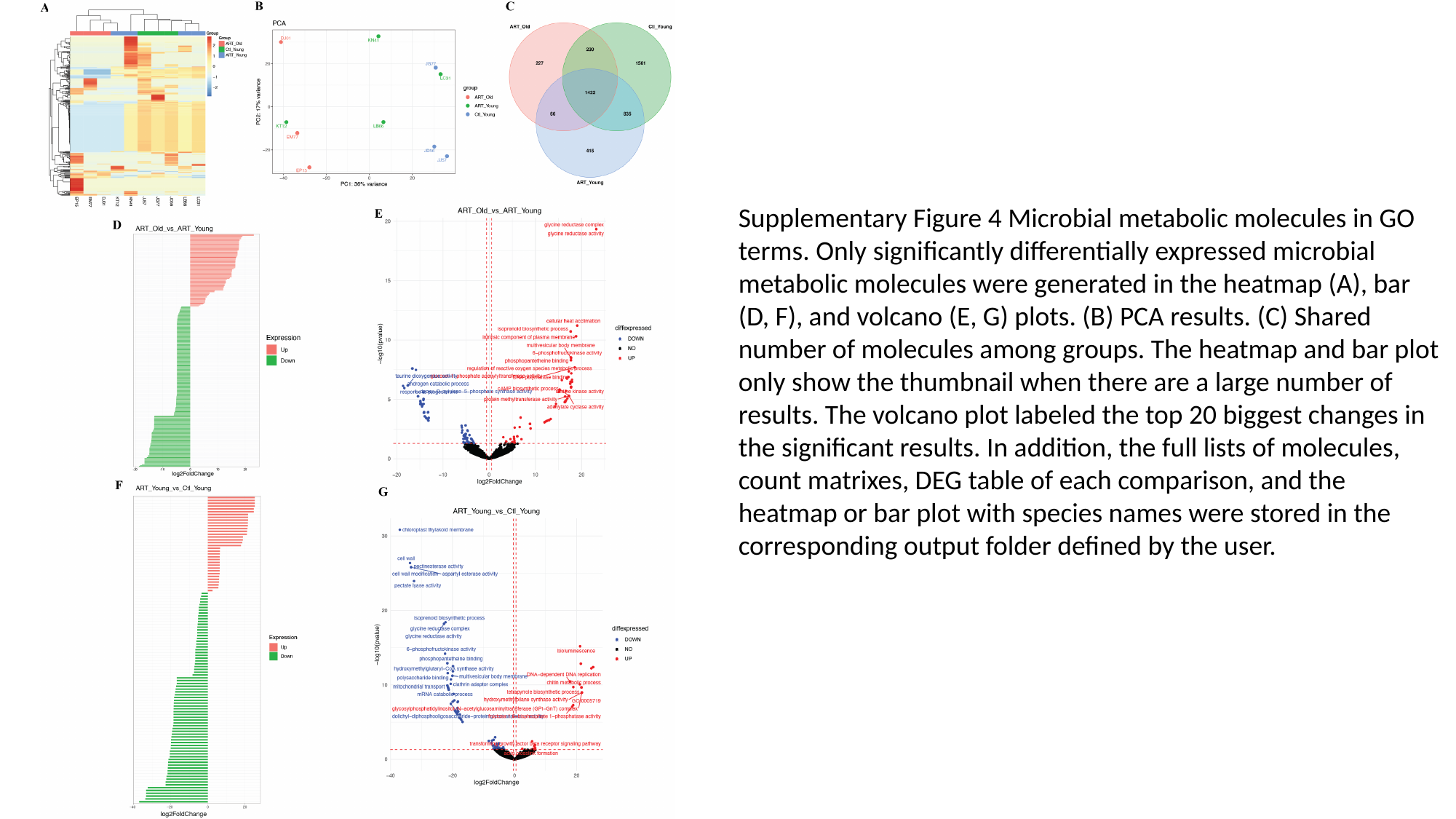

Supplementary Figure 4 Microbial metabolic molecules in GO terms. Only significantly differentially expressed microbial metabolic molecules were generated in the heatmap (A), bar (D, F), and volcano (E, G) plots. (B) PCA results. (C) Shared number of molecules among groups. The heatmap and bar plot only show the thumbnail when there are a large number of results. The volcano plot labeled the top 20 biggest changes in the significant results. In addition, the full lists of molecules, count matrixes, DEG table of each comparison, and the heatmap or bar plot with species names were stored in the corresponding output folder defined by the user.

### Slide 2
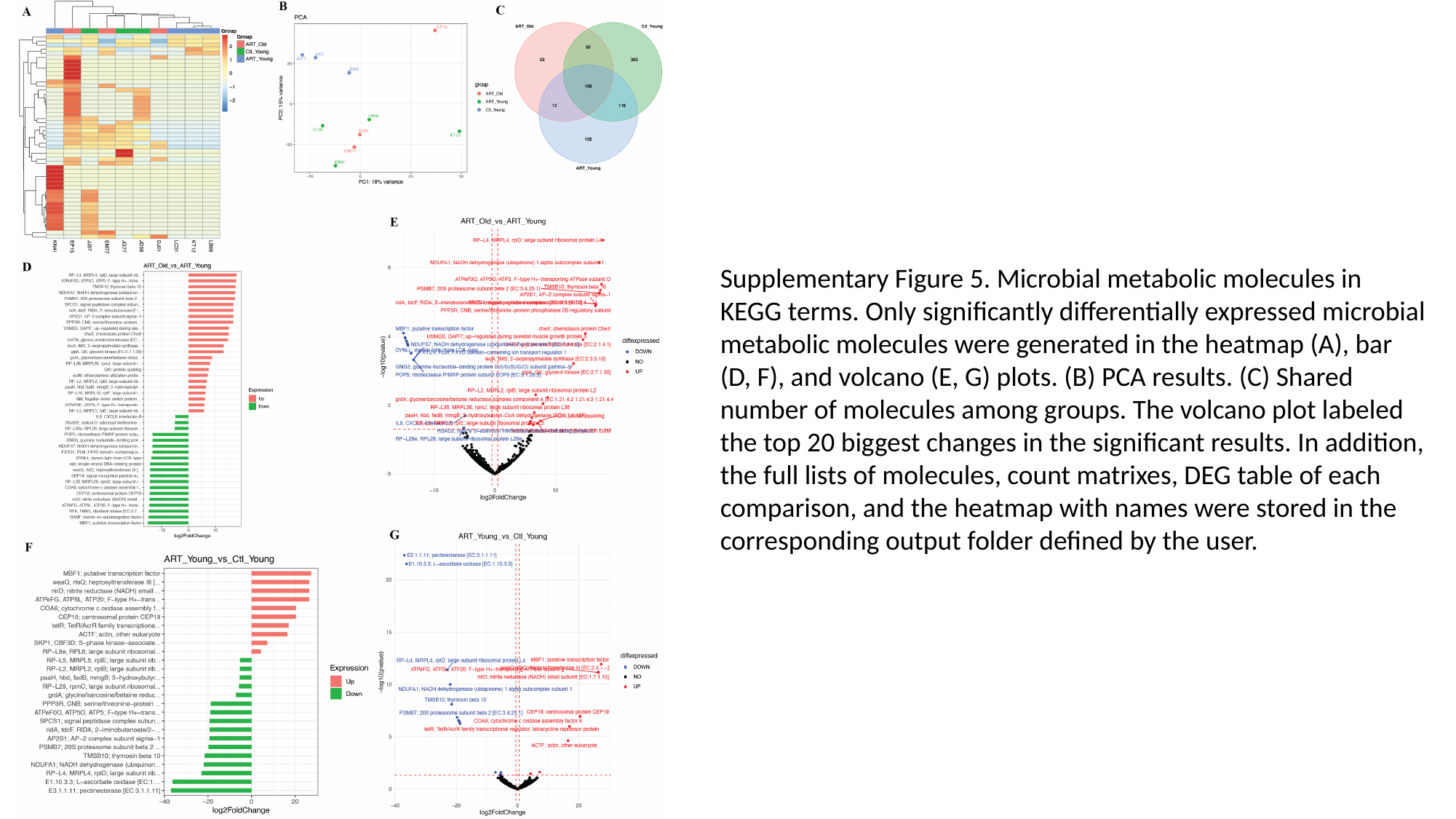

Supplementary Figure 5. Microbial metabolic molecules in KEGG terms. Only significantly differentially expressed microbial metabolic molecules were generated in the heatmap (A), bar (D, F), and volcano (E, G) plots. (B) PCA results. (C) Shared number of molecules among groups. The volcano plot labeled the top 20 biggest changes in the significant results. In addition, the full lists of molecules, count matrixes, DEG table of each comparison, and the heatmap with names were stored in the corresponding output folder defined by the user.
