## Supplementary Figure 6 for "MTD: a unique pipeline for host and meta-transcriptome joint and integrative analyses of RNA-seq data"

### Slide 1
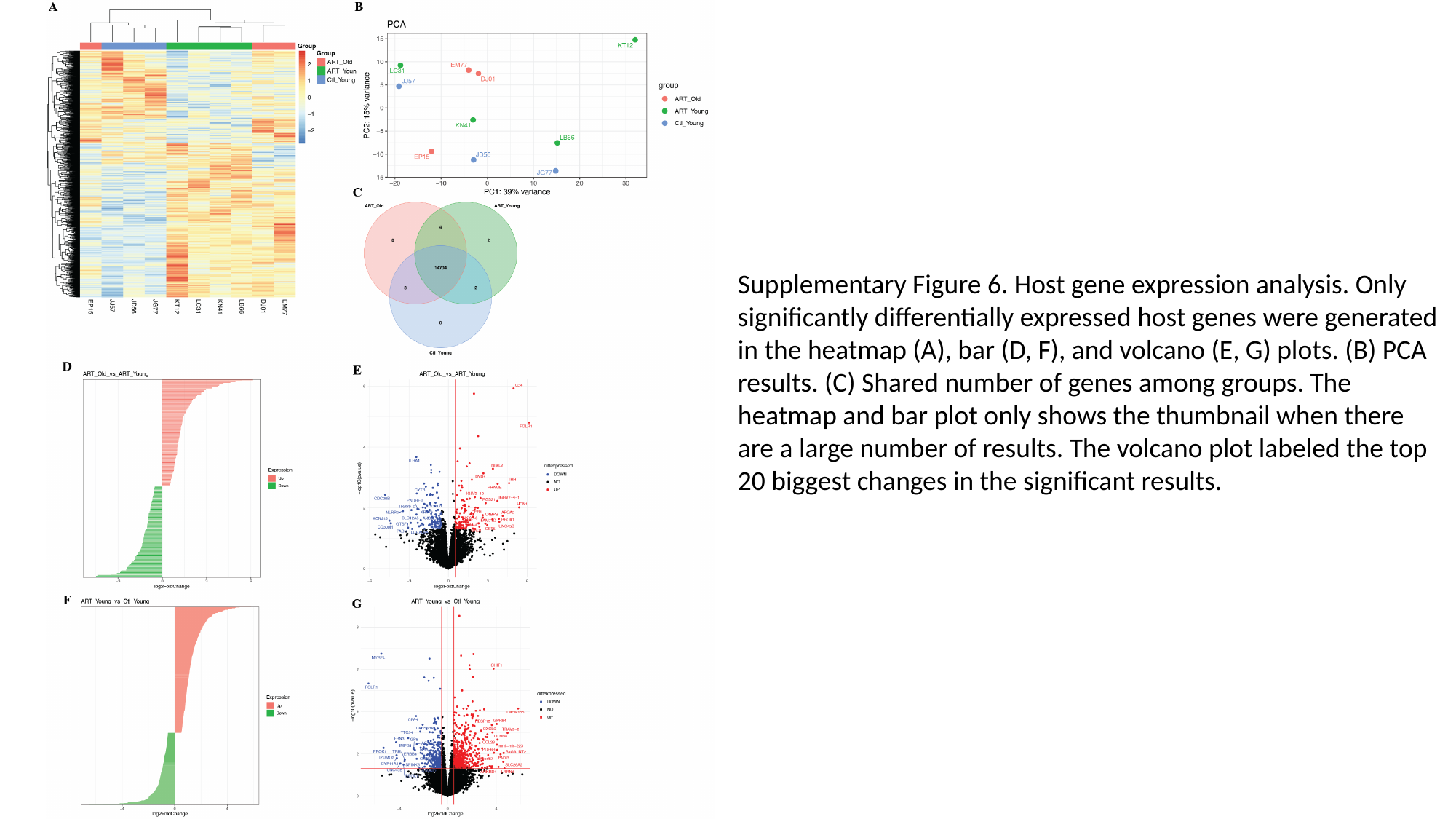

Supplementary Figure 6. Host gene expression analysis. Only significantly differentially expressed host genes were generated in the heatmap (A), bar (D, F), and volcano (E, G) plots. (B) PCA results. (C) Shared number of genes among groups. The heatmap and bar plot only shows the thumbnail when there are a large number of results. The volcano plot labeled the top 20 biggest changes in the significant results.
